# Pervasive effects of *Wolbachia* on host activity

**DOI:** 10.1101/2021.02.24.432688

**Authors:** Michael T.J. Hague, H. Arthur Woods, Brandon S. Cooper

**Affiliations:** Division of Biological Sciences, University of Montana, 32 Campus Dr. Missoula, MT 59812

## Abstract

Heritable symbionts have diverse effects on the physiology, reproduction, and fitness of their hosts. Maternally transmitted *Wolbachia* are one of the most common endosymbionts in nature, infecting about half of all insect species. We test the hypothesis that *Wolbachia* alter host behavior by assessing the effects of 14 different *Wolbachia* strains on the locomotor activity of nine *Drosophila* host species. We find that *Wolbachia* alter the activity of six different host genotypes, including all hosts in our assay infected with *w*Ri-like *Wolbachia* strains (*w*Ri, *w*Suz, *w*Aur), which have rapidly spread among *Drosophila* species in only the last 13,000 years. While *Wolbachia* effects on host activity were common, the direction of these effects varied unpredictability and sometimes depended on host sex. We hypothesize that the prominent effects of *w*Ri-like *Wolbachia* may be explained by patterns of *Wolbachia* titer and localization within host somatic tissues, particularly in the central nervous system. Our findings support the view that *Wolbachia* have wide-ranging effects on host behavior. The fitness consequences of these behavioral modifications are important for understanding the evolution of host-symbiont interactions, including how *Wolbachia* spread within host populations.

## INTRODUCTION

Insects harbor microorganisms that have wide-ranging effects on their performance and fitness [1–3], including manipulations to reproduction [4–7], provisioning of nutrients [1,8,9], modifications of thermotolerance [10,11], and defense against pathogens [12–15]. Microbes may also alter host behavior [16–21]. In extreme instances, parasitic microbes can induce behaviors that increase the likelihood of transmission—for example, by directing hosts to habitats that promote transmission [22–28]. Infected hosts may also change their own behavior as an immune strategy against infection, including seeking warm temperatures to induce a “behavioral fever” [29,30] or reducing activity and increasing sleep time [19,31–34]. Such behavioral modifications have important implications for microbe spread and host fitness.

Maternally transmitted *Wolbachia* are the most common endosymbionts in nature, infecting many arthropods [5,35,36] and two distantly related groups of nematodes [37]. Discordant *Wolbachia* and host phylogenies indicate that many hosts have recently acquired *Wolbachia* via introgressive and horizontal transfer [38–43]. *Wolbachia* are primarily transmitted vertically by female hosts, so natural selection favors beneficial effects on host fitness that promote spread [44–47]. Maternal transmission occurs via the host reproductive system, but *Wolbachia* are also found in host somatic tissues, including nervous, digestive, and metabolic tissues [48–51]. Still, the behavioral and physiological consequences of somatic infections are poorly understood [19,51].

Prior work indicates *Wolbachia* influence several host behaviors [19,52,53], including sleep [54–56] and temperature preference [20,57,58]. We broadly test for *Wolbachia* effects on the locomotor activity of *Drosophila* hosts infected with A-group *Wolbachia* (*N* = 11), B-group *Wolbachia* (*N* = 1), and an A- and B-group co-infection (*N* = 1). Our analysis includes two prominent A-group clades that recently spread among *Drosophila*: *w*Mel-like *Wolbachia* (*w*MelCS, two *w*Mel variants, *w*Yak, *w*San, and *w*Tei) and *w*Ri-like *Wolbachia* (*w*Ri, *w*Suz, and *w*Aur) [42,43]. We find that *Wolbachia* effects on host activity are common, particularly for *w*Ri-like *Wolbachia*, a “super-spreader” strain that rapidly spread among *Drosophila* species in the last ~13,000 years [42].

## METHODS

### Fly lines

We evaluated 13 different *Wolbachia-infected* host genotypes (Figure 1, Table S1), consisting of nine *Drosophila* species infected with 14 different A- and B-group *Wolbachia* that diverged up to 46 million years ago [59]. For two of the host species, *D. melanogaster* and *D. simulans*, we tested multiple *Wolbachia-infected* genotypes. This included a *D. simulans* host co-infected with A-group *w*Ha and B-group *w*No [60–63]. We used tetracycline treatment as previously described [20] to generate uninfected genotypes to pair with each infected genotype, while taking care to avoid detrimental effects of the antibiotic treatment on mitochondrial function [64] (see Supplemental Methods).

**Figure 1.**
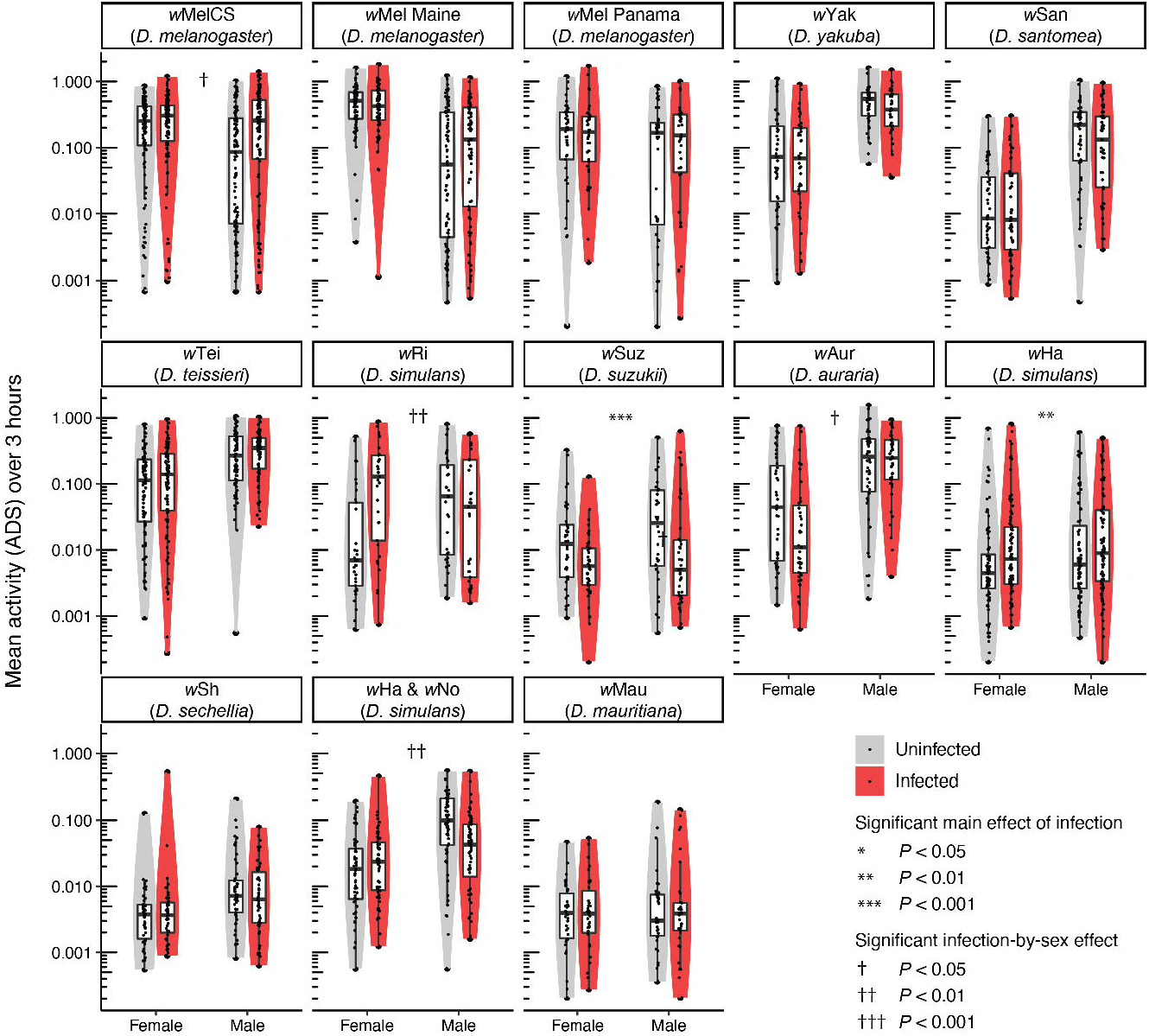
Activity of uninfected and infected flies for each sex of each genotype. Activity is measured as mean absolute distance sums (ADS). Significance was evaluated using linear models (Tables S2 and S3).

### Host locomotor activity assays

We reared flies at 25°C under a 12-h light: 12-h dark cycle (Percival model I-36LL) on a standard food diet [20]. Each day, we collected a batch of female and male virgins for one pair of uninfected and infected genotypes. The four treatment groups (uninfected females, infected females, uninfected males, and infected males) were maintained in isolation until they were 3 to 5 days old. We then measured the locomotor activity of the batch of flies using a 16-chamber flow-through respirometry and data acquisition system (MAVEn, Sable Systems International). The MAVEn has 16 2.4 ml volume polycarbonate animal chambers and an activity board that uses infrared light (invisible to flies) to monitor animal activity in each chamber, sampled at 1 Hz (Figure S1). Individual flies were aspirated into a randomly assigned chamber and allowed to adjust to the new environment for 0.5 hours. Activity measurements were then recorded over a 3hour period between the hours of 0900 and 1600.

The raw outputs from the activity sensors were transformed into the activity index absolute distance sums (ADS). We calculated ADS by first calculating the cumulative sum of the absolute difference between consecutive activity readings, and then calculating the slope of cumulative activity vs. time [65,66]. We used mean ADS over the 3-hour period as our estimate of locomotor activity for each fly; however, our analyses were robust regardless of how we quantified activity (see Supplemental Methods). We found that the mean ADS activity data required a transformation for statistical analysis; however, a single data transformation was not suitable for all host species. We used a log transformation of mean ADS for *D. simulans, D. suzukii, D. auraria, D. mauritiana*, and *D. sechellia*, and a square root transformation for *D. melanogaster, D. yakuba, D. santomea*, and *D. teissieri.* We present a full statistical analysis of all datasets in Tables S2 and S3, respectively.

We used the log- and square root-transformed mean ADS data as dependent variables in linear models. We included infection status, sex, and an infection-by-sex interaction effect as independent variables, as well as additional independent variables to account for other potential sources of activity variation: randomly assigned animal chamber (1-16), experimental start time, mean water vapor (ppt), mean relative humidity (%), mean temperature (°C), and mean light intensity (lux) [65,66]. We evaluated the significance of individual effects using *F* tests and type III sum of squares using the “Anova” function in the *car* R package [67,68].

### Wolbachia phylogenomic analysis

We used publicly available *Wolbachia* genome assemblies [20,42,43,59,69–71], and new Illumina sequencing, to generate a Bayesian phylogram [20] (see Supplemental Methods). *Wolbachia* effects on host activity were especially common for *w*Ri-like *Wolbachia*, so we used the phylogram to test whether *Wolbachia* effects on hosts exhibit phylogenetic signal. First, we treated *Wolbachia* effects on host locomotor activity as a binary trait and tested for phylogenetic signal using the *D* statistic [72], implemented in the *caper* R package [73]. Second, we treated *Wolbachia* effects on activity as a continuous trait and tested for phylogenetic signal using Pagel’s lambda (λ) [74]. Here, we analyzed each sex separately, because we found significant infection-by-sex interaction effects on activity (Tables S2 and S3). For each sex, we extracted the least-square (LS) mean ADS for infected and uninfected flies from the linear models (Tables S2 and S3), and used the change in LS mean activity as a continuous character to calculate the maximum likelihood value of Pagel’s λ [74,75]. We used a likelihood ratio test to compare our fitted value of λ to a model assuming no phylogenetic signal (λ = 0) using the “phylosig” function in the R package *phytools* [76].

## RESULTS

### Wolbachia infections modify host locomotor activity

We assayed the locomotor activity of 3,104 flies (Figure 1). *Wolbachia* had a significant effect on the activity of six host genotypes, including hosts infected with both A- and B-group *Wolbachia.* Interestingly, the direction of *Wolbachia* effects on host activity varied by genotype and sex (Figure 2). We found a significant *Wolbachia* infection-by-sex interaction effect for the *w*MelCS-*D. melanogaster* genotype that increased male activity (*F* = 4.566, *P* = 0.033; Table S3). We also found a significant infection-by-sex effect for the *w*Ri-*D. simulans* genotype, but *Wolbachia* increased female activity (*F* = 8.150, *P* = 0.005; Table S2). The two other closely related *w*Ri-like *Wolbachia, w*Suz and *w*Aur, also had significant effects on host activity. The *w*Suz-*D. suzukii* genotype had a significant main effect of *Wolbachia* that reduced host activity (*F* = 11.311, *P* < 0.001; Table S2), and the *w*Aur-*D. auraria* genotype had a significant infection-by-sex interaction that reduced female activity (*F* = 6.584, *P* = 0.011; Table S2). The *w*Ha-*D. simulans* genotype had a significant main effect of *Wolbachia* that increased host activity (*F* = 7.764, *P* = 0.006; Table S2). Lastly, we found the *w*Ha-*w*No co-infected *D. simulans* genotype had a significant infection-by-sex interaction effect that reduced male activity (*F* = 7.076, *P* = 0.008; Table S2). Because this genotype is co-infected, we do not know the relative contributions of *w*Ha and *w*No to variation in host activity. See the Supplemental Results for a discussion of how other variables contributed to variation in locomotor activity.

**Figure 2.**
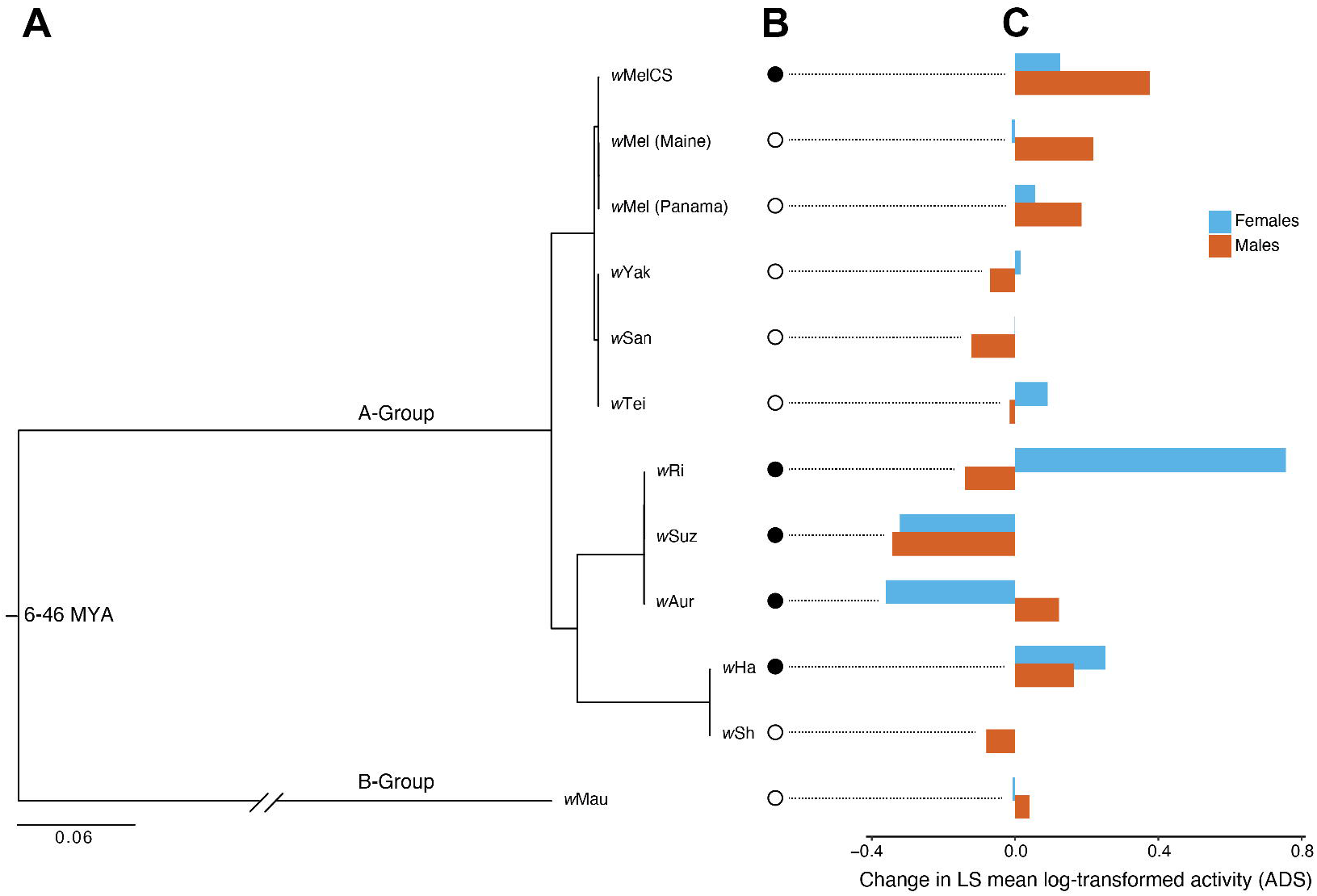
(**A**) Estimated Bayesian phylogram for A- and B-group *Wolbachia* strains. The divergence estimate for A- and B-groups is superimposed from Meany et al. [59]. All nodes have Bayesian posterior probabilities of 1. (**B**) *Wolbachia* effects on host activity scored as a binary trait: *Wolbachia* significantly altered host activity (black circle) or had no effect (white circle). (**C**) *Wolbachia* effects on activity scored as a continuous trait: the change in least-square (LS) mean log-transformed activity (ADS) for each sex. LS means were generated from linear models (Table S2). LS mean square root-transformed ADS data are shown in Figure S3.

### Limited evidence for phylogenetic signal

We estimated a Bayesian phylogram of A- and B-group *Wolbachia* using 211 single-copy genes of identical length in all *Wolbachia* genomes, spanning 178,569 bp (Figure 2). We then tested whether closely related *Wolbachia* have similar effects on host activity. When treating *Wolbachia* effects on activity as a binary trait, our estimate of *D* = 0.322 was low, but not statistically different from a model of *D* = 1 assuming phylogenetic randomness (*P* = 0.101) or a model of *D* = 0 with strong phylogenetic signal (*P* = 0.198). Simulations of similar phylogenies with an increasing number of *Wolbachia* strains suggest that at least *N* = 50 strains are required to differentiate our estimated value of *D* = 0.322 from a model of phylogenetic randomness (*D* = 1) (Figures S4 and S5). Thus, *Wolbachia* effects on host activity may exhibit phylogenetic signal, but many more *Wolbachia* strains are required to test this hypothesis. Unfortunately, *N* = 50 strains are not presently available in culture. We also treated *Wolbachia* changes to host activity as a continuous trait; however, we found that maximum likelihood fitted λ values were extremely low, indicative of no phylogenetic signal. λ values generated from the LS mean log-transformed ADS data were not statistically different from zero for females (λ < 0.001, *P* = 1) or males (λ < 0.001, *P* = 1). This was also true when we repeated the analyses for the LS mean square root-transformed ADS data for females (λ < 0.001, *P* = 1) and males (λ < 0.001, *P* = 1).

## DISCUSSION

Our analyses suggest that *Wolbachia* commonly alter host locomotor activity, which may affect host fitness. Locomotion is a basic host activity underlying many ecologically important behaviors, including foraging, thermoregulation, and mate seeking. In combination with our recent work demonstrating pervasive effects of A- and B-group *Wolbachia* on host temperature preference [20], we posit that *Wolbachia* infections may often alter host behavior.

The *w*Ri-like *Wolbachia* strains in our study (*w*Ri, *w*Suz, and *w*Aur) consistently altered host activity. We found a low, but non-significant *D* value of 0.322, suggesting effects on host activity may exhibit phylogenetic signal; although, an excessive number of *Wolbachia* strains are required to test this hypothesis. Our findings are consistent with prior experiments demonstrating that *w*Ri increased female *D. simulans* activity in response to olfactory cues [77,78]. We hypothesize that the prominence of *w*Ri-like *Wolbachia* effects on host activity relative to other strains may be due to variation in *Wolbachia* tissue localization [49,52]. *w*Ri occurs at high titer in adult *D. simulans* brains and localizes to specific regions, whereas *w*Mel shows a relatively even distribution in *D. melanogaster* [52]. *w*Ri also occurs at higher titer in the ventral nerve cord, which is a major neural circuit center for motor activities such as walking [52,79–81]. Future experiments should compare *Wolbachia* titer and localization in adult brains for *w*Ri-like variants and strains that do not alter locomotor activity.

We also found considerable variation in the direction and sex-bias of *Wolbachia* effects on locomotor activity (Figure 2). *Wolbachia* decreased activity for *w*Suz, *w*Aur, and the *w*No-*w*Ha co-infection, whereas *w*MelCS, *w*Ri, and *w*Ha increased activity. These effects were female-biased for *w*Ri and *w*Aur, but male-biased for *w*MelCS and *w*No-*w*Ha. This variation had no relationship to the *Wolbachia* phylogeny, because we found no evidence for phylogenetic signal when measuring *Wolbachia* effects on females and males as a continuous trait (λ < 0.001). Specific *Wolbachia* effects on host activity may depend on interactions with the host background. For example, our work and others’ suggests that identical *w*MelCS variants have different effects on *D. melanogaster* temperature preference depending on the host background [20,57,58]. Host genomes also modify *Wolbachia* titer [82], *Wolbachia* maternal transmission [83], components of host fitness [84–87], and the strength of cytoplasmic incompatibility [88–90].

Changes to host activity could underlie *Wolbachia*-induced behaviors that promote infection spread. For example, *w*Mel-infected *D. melanogaster* have higher field recapture rates than uninfected flies [91], and long distance dispersal of the spider *Erigone atra* is altered by *Rickettsia,* an endosymbiont closely related to *Wolbachia* [92]. Our own work suggests *Wolbachia* may alter host temperature preference to promote *Wolbachia* replication within host bodies [20]. Other experiments suggest that *w*Mel and *w*Ri may influence male mating rate [93,94]. Alternatively, hosts may be modifying their own behavior as a response to *Wolbachia* infection. Several studies indicate that *w*Mel alters circadian activity and sleep patterns of *D. melanogaster* [52,54–56]. For example, Bi et al. [55] report that *w*Mel increases sleep time, which could represent a host immune response to infection [19]. Ultimately, these effects on host behavior factor into how *Wolbachia* influence host fitness, which determines the spread and persistence of *Wolbachia* in host populations [4,95–98]. Because locomotor activity is such a fundamental host behavior, our results suggest *Wolbachia* may have complex and variable effects on many components of host fitness.

## ACKNOWLEDEMENTS

We thank TBW for lab assistance and WRC for help with bioinformatic analyses. The Cooper lab group and MT provided valuable feedback that improved the manuscript. This work was supported by NIGMS of the NIH under award R35GM124701 to BSC.

## DATA AVAILABILITY

All data and code are available on Dryad (doi:10.5061/dryad.6t1g1jwxv; temporary URL: https://datadryad.org/stash/share/D4Yi8zYFchXE-LIFgTc6S91s2Nl5Afcwu4-FkxJZPaA). Genome assemblies will be deposited on GenBank upon acceptance.

