## Supplemental Materials for "Pervasive effects of *Wolbachia* on host activity"

### SUPPLEMENTAL METHODS

*Tetracycline treatment*

We generated *Wolbachia*-uninfected genotypes by treating the infected lines with 0.03% tetracycline for four generations. After the fourth generation, we used quantitative PCR (qPCR) on 5 females homogenized together to confirm that flies were cleared of *Wolbachia*. Genomic DNA was extracted using a DNeasy Blood & Tissue kit (Qiagen). We used a Stratagene Mx3000P (Agilent Technologies) to amplify the *Wolbachia*-specific locus *ftsZ* using forward (5'-GCAGCCAATAGAGTGCGTG) and reverse (5'-TTCCCTCCATCGCTTGATCA) primers. Efficiency curves were generated to confirm the primers had adequate efficiency. We used the following cycling conditions: 50°C for 2 minutes, 95°C for 2 minutes, and then 40 cycles, with one cycle consisting of 95°C for 15 seconds and 60°C for 1 minute. We ran three technical replicates for each sample to confirm the removal of *Wolbachia*. We also amplified the *Drosophila*-specific *UCE* locus using forward (5'- GAAATAAACGCAAGCGCCATC) and reverse (5'- GCAAGCGCGTCTCAAGC) primers that served as a positive control. The qPCR primers were kindly provided by Dylan Shropshire. Finally, we reconstituted the gut microbiome of the tetracycline-cleared flies by rearing them on food where infected males of the same genotype had fed and defecated for the prior 48 hours. Tetracycline-cleared flies were given at least three more generations before we conducted experiments to avoid detrimental effects of the antibiotic treatment on mitochondrial function (Ballard and Melvin 2007). An alternative approach to generate pairs of *Wolbachia*-infected and uninfected genotypes would be to reciprocally introgress their cytoplasms; however, this approach fails to control mitochondria that may affect host energetics and activity. Thus, we strategically chose to generate paired uninfected genotypes using tetracycline treatment of infected genotypes.

*Host locomotor activity assays*

To ensure that our analyses were robust regardless of how we quantified activity over the 3-hour period, we repeated our statistical analyses using a second index of activity that is analogous to measures of locomotor activity from *Drosophila* Activity Monitoring Systems (DAMS) (Chiu et al. 2010; Pfeiffenberger et al. 2010). Here, we counted the number of "activity events" or the cumulative number of seconds that each fly triggered the infrared activity monitor over the 3-hour period (Figure S2). These count data were highly correlated with our estimates of mean ADS (Pearson's $r = 0.952$, $P < 0.001$), and our statistical analysis of the log- and square root-transformed count data (Tables S4 and S5, respectively) produced highly similar results to the respective log- and square root-transformed mean ADS analyses (Tables S2 and S3, respectively). Thus, only results from the mean ADS analysis are presented in the main text.

*Wolbachia sequencing and phylogenomic analysis*

We obtained *Wolbachia* sequences from publicly available genome assemblies, which included *w*Ri (Klasson et al. 2009), *w*Ha (Ellegaard et al. 2013), *w*Mau (Meany et al. 2019), *w*Yak, *w*Tei, *w*San (Cooper et al. 2019), *w*Suz (Siozios et al. 2013), and *w*Sh (Hague et al. 2020). Because a reference genome is not currently available for *w*Aur, we obtained raw Illumina reads from a *w*Aur-infected *D. auraria* individual from a previously published dataset (Accession PRJNA432099) (Turelli et al. 2018). We obtained *w*Mel assemblies for our *Canton-S Berkeley* and *PC75 D. melanogaster* genotypes from our previous study using the same strains (Accession PRJNA65830) (Hague et al. 2020). We sequenced the genome of the other *w*Mel genotype included in this study (*FFD25*) using previously described methods (Hague et al. 2020). Because we could not attribute effects on host activity to an individual *Wolbachia* strain for our *w*Ha-*w*No co-infected *D. simulans* genotype (*sim*198), we did not consider *w*No in our phylogenomic analysis and we used the singly infected *w*Ha genotype (*Car5*) to estimate *w*Ha effects on activity in our tests for phylogenetic signal.

Raw Illumina reads were trimmed using Sickle version 1.33 (Joshi et al. 2011) and assembled using ABySS version 2.0.2 (Jackman et al. 2017). *K* values of 71, 81, and 91 were used, and scaffolds with the best nucleotide BLAST matches to known *Wolbachia* sequences with *E*-values less than $10^{-10}$ were extracted as the draft *Wolbachia* assemblies. For each genotype, we chose the assembly with the highest N50 and the fewest scaffolds (Table S6). The *w*Aur genome, *w*Mel from *FFD25*, and the ten previously published genomes were annotated using Prokka version 1.11, which identifies homologs to known bacterial genes (Seemann 2014). To avoid pseudogenes and paralogs, we only used genes present in a single copy with no alignment gaps in all of the genome sequences. Genes were identified as single copy if they uniquely matched a bacterial reference gene identified by Prokka. By requiring all homologs to have identical length in all of the *Wolbachia* genomes, we removed all loci with indels. A total of 211 genes totaling 178,569 bp met these criteria. We then estimated a Bayesian phylogram using

RevBayes 1.0.8 under the GTR + Γ + I model partitioned by codon position (Höhna et al. 2016). Four independent runs were performed, which all converged on the same topology. All nodes were supported with Bayesian posterior probabilities of 1.

### SUPPLEMENTAL RESULTS

Four of the genotypes (*w*MelCS, *w*Ri, *w*Aur, and *w*No-*w*Ha) had significant infection-by-sex interaction effects on host locomotor activity, indicating that sex is also an important determinant of activity. In addition, we found that sex had a significant main effect on locomotor activity for six other genotypes: *w*Mel-*D. melanogaster* from Maine, *w*Yak-*D. yakuba*, *w*San-*D. santomea*, *w*Tei-*D. teissieri*, *w*Ha-*D. simulans*, and *w*Sh-*D. sechellia*. Females were more active than males for the *w*Mel-*D. melanogaster* Maine genotype ($F$ = 153.961, $P$ < 0.001; Table S3). In the *D. yakuba* clade of hosts, males were more active than females for the *w*Yak-*D. yakuba* genotype ($F$ = 67.904, $P$ < 0.001), the *w*San-*D. santomea* genotype ($F$ = 114.888, $P$ < 0.001), and the *w*Tei-*D. teissieri* genotype ($F$ = 93.461, $P$ < 0.001; Table S3). Finally, males were more active than females for the *w*Ha-*D. simulans* genotype ($F$ = 4.737, $P$ = 0.03) and the *w*Sh-*D. sechellia* genotype ($F$ = 26.288, $P$ < 0.001; Table S2).

In addition to *Wolbachia* and sex, we found other significant effects on locomotor activity. Experimental start times later in the day reduced activity of the *w*MelCS-*D. melanogaster* genotype ($F$ = 6.642, $P$ = 0.01) and the *w*Mel-*D. melanogaster* genotypes from Maine ($F$ = 18.839, $P$ <0.001) and Panama City ($F$ = 13.984, $P$ < 0.001; Table S3), but increased activity of the *w*Suz-*D. suzukii* genotype ($F$ = 6.756, $P$ = 0.01; Table S2). Increased water vapor (ppt) caused a decline in activity for the *w*MelCS-*D. melanogaster* genotype ($F$ = 12.06, $P$ < 0.001), the *w*San-*D. santomea* genotype ($F$ = 11.196, $P$ < 0.001), the *w*Tei-*D. teissieri* genotype ($F$ = 7.491, $P$ = 0.007; Table S3), and the *w*Sh-*D. sechellia* genotype ($F$ = 6.853, $P$ = 0.01), but caused an increase in activity for the *w*Aur-*D. auraria* genotype ($F$ = 16.281, $P$ < 0.001; Table S2). Increased relative humidity (%) resulted in a decline in activity for the *w*Mel-*D. melanogaster* genotype from Maine ($F$ = 4.462, $P$ = 0.035; Table S3) and the *w*Mau-*D. maurtiana* genotype ($F$ = 5.462, $P$ = 0.021), but an increase in activity for the *w*Suz-*D. suzukii* genotype ($F$ = 4.749, $P$ =0.031), the *w*Aur-*D. auraria* genotype ($F$ = 9.767, $P$ = 0.002), and the co-infected *w*Ha-*w*No-*D. simulans* genotype ($F$ = 11.104, $P$ = 0.001; Table S2). Increased ambient temperature (°C) caused an increase in activity for the *w*Mel-*D. melanogaster* genotype from Panama City ($F = 3.959$, $P = 0.049$), the *w*Yak-*D. yakuba* genotype ($F = 5.032$, $P = 0.026$), the *w*San-*D. santomea* genotype ($F = 24.25$, $P < 0.001$), and the *w*Tei-*D. teissieri* genotype ($F = 3.864$, $P = 0.05$; Table S3), but caused a decline in activity for the *w*Suz-*D. suzukii* genotype ($F = 9.125$, $P = 0.003$; Table S2). Finally, increased ambient light intensity (lux) decreased activity of the *w*Tei-*D. teissieri* genotype ($F = 6.912$, $P = 0.009$; Table S3), but increased activity of the *w*Suz-*D. suzukii* genotype ($F = 10.884$, $P = 0.001$) and the *w*Aur-*D. auraria* genotype ($F = 19.044$, $P < 0.001$; Table S2).

### SUPPLEMENTAL TABLES

**Supplemental Table S1.** Genotype IDs for different *Wolbachia*-infected host species used in this study. Genotype IDs marked with an asterisk were obtained from our prior study on *Wolbachia* effects on host temperature preference (Hague et al. 2020). With the exception of the *Canton-S Berkeley*, all *Wolbachia*-infected isofemale lines were formed by naturally sampling single gravid females from the field and placing them individually into vials.

| Species | *Wolbachia* | Genotype ID | Collection Location |
|---|---|---|---|
| *D. melanogaster* | *w*MelCS | *Canton-S Berkeley** | Canton, Ohio, USA |
| *D. melanogaster* | *w*Mel | *FFD25* | Maine, USA |
| *D. melanogaster* | *w*Mel | *PC75** | Panama City, Panama |
| *D. yakuba* | *w*Yak | *B13L5** | Bioko, Equatorial Guinea |
| *D. santomea* | *w*San | *S09L4* | São Tomé, São Tomé and Príncipe |
| *D. teissieri* | *w*Tei | *B13L11** | Bioko, Equatorial Guinea |
| *D. simulans* | *w*Ri | *Riv84** | Riverside, California, USA |
| *D. simulans* | *w*Ha | *Car5** | Hawaii, USA |
| *D. simulans* | *w*Ha & *w*No | *sim198* | New Caledonia |
| *D. suzukii* | *w*Suz | *S278* | California, USA |
| *D. auraria* | *w*Aur | *L2* | Japan |
| *D. sechellia* | *w*Sh | *PmuseumbananaI** | Praslin, Seychelles |
| *D. mauritiana* | *w*Mau | *mauR31** | Mauritius |

**Supplemental Table S2.** Results from linear models of log-transformed mean absolute distance sums (ADS). Leverage and distribution analyses revealed that the mean ADS data for *D. simulans*, *D. suzukii, D. auraria*, *D. mauritiana*, and *D. sechellia* required a log transformation. Significant effects identified by *F* tests at the *P* = 0.05 threshold are marked with an asterisk.

| | ***w*MelCS** | | ***w*Mel (Maine)** | | ***w*Mel (Panama)** | | ***w*Yak** | | ***w*San** | |
|---|---|---|---|---|---|---|---|---|---|---|
| **Explanatory variable** | ***F*** | ***P* value** | ***F*** | ***P* value** | ***F*** | ***P* value** | ***F*** | ***P* value** | ***F*** | ***P* value** |
| Infection Status | 11.484 | <0.001* | 1.869 | 0.173 | 0.915 | 0.341 | 0.084 | 0.773 | 0.445 | 0.505 |
| Sex | 13.029 | <0.001* | 133.505 | <0.001* | 3.308 | 0.071 | 68.459 | <0.001* | 111.668 | <0.001* |
| Chamber | 1.541 | 0.093 | 2.635 | 0.001* | 1.427 | 0.148 | 1.817 | 0.042* | 1.361 | 0.176 |
| Start Time | 4.990 | 0.026* | 20.769 | <0.001* | 10.205 | 0.002* | 0.016 | 0.899 | 0.156 | 0.693 |
| Water Vapor Pressure | 7.886 | 0.005* | 0.189 | 0.664 | 0.875 | 0.351 | 0.138 | 0.711 | 13.645 | <0.001* |
| Relative Humidity | 2.305 | 0.130 | 4.175 | 0.042 | 0.372 | 0.543 | 0.879 | 0.350 | 1.489 | 0.224 |
| Temperature | 1.305 | 0.254 | 1.600 | 0.207 | 5.050 | 0.026* | 3.286 | 0.072 | 26.711 | <0.001* |
| Light Intensity | 0.847 | 0.358 | 0.618 | 0.432 | 0.058 | 0.810 | 0.014 | 0.908 | 0.262 | 0.609 |
| Infection-by-Sex | 2.854 | 0.092 | 2.167 | 0.142 | 0.254 | 0.615 | 0.222 | 0.638 | 0.424 | 0.516 |
| Sample Size | 464 | | 341 | | 160 | | 164 | | 209 | |

| | ***w*Tei** | | ***w*Ri** | | ***w*Suz** | | ***w*Aur** | | ***w*Ha** | |
|---|---|---|---|---|---|---|---|---|---|---|
| **Explanatory variable** | ***F*** | ***P* value** | ***F*** | ***P* value** | ***F*** | ***P* value** | ***F*** | ***P* value** | ***F*** | ***P* value** |
| Infection Status | 0.377 | 0.539 | 5.091 | 0.026* | 11.311 | <0.001* | 1.484 | 0.225 | 7.764 | 0.006* |
| Sex | 78.745 | <0.001* | 0.384 | 0.537 | 3.493 | 0.064 | 69.740 | <0.001* | 4.737 | 0.030* |
| Chamber | 3.184 | <0.001* | 0.664 | 0.814 | 1.580 | 0.092 | 2.411 | 0.004* | 3.420 | <0.001* |
| Start Time | 4.897 | 0.028* | 3.868 | 0.052 | 6.756 | 0.010* | 0.609 | 0.436 | 3.310 | 0.070 |
| Water Vapor Pressure | 11.127 | <0.001* | 0.001 | 0.979 | 1.257 | 0.264 | 16.281 | <0.001* | 0.536 | 0.465 |
| Relative Humidity | 0.018 | 0.893 | 0.450 | 0.504 | 4.749 | 0.031* | 9.767 | 0.002* | 0.251 | 0.617 |
| Temperature | 0.939 | 0.333 | 0.775 | 0.381 | 9.125 | 0.003* | 0.000 | 0.988 | 1.303 | 0.254 |
| Light Intensity | 3.778 | 0.053 | 2.348 | 0.128 | 10.884 | 0.001* | 19.044 | <0.001* | 0.828 | 0.363 |
| Infection-by-Sex | 0.672 | 0.413 | 8.150 | 0.005* | 0.012 | 0.914 | 6.584 | 0.011* | 0.349 | 0.555 |
| Sample Size | 347 | | 131 | | 163 | | 181 | | 353 | |

| | ***w*Sh** | | **wHa & wNo** | | ***w*Mau** | |
|---|---|---|---|---|---|---|
| **Explanatory variable** | ***F*** | ***P* value** | ***F*** | ***P* value** | ***F*** | ***P* value** |
| Infection Status | 0.380 | 0.538 | 3.939 | 0.048 | 0.040 | 0.842 |
| Sex | 26.288 | <0.001* | 32.081 | <0.001* | 0.005 | 0.947 |
| Chamber | 2.452 | 0.003* | 1.127 | 0.335 | 3.682 | <0.001* |
| Start Time | 1.428 | 0.234 | 3.329 | 0.069 | 1.257 | 0.264 |
| Water Vapor Pressure | 6.853 | 0.010* | 2.464 | 0.118 | 0.277 | 0.599 |
| Relative Humidity | 0.311 | 0.578 | 11.104 | 0.001* | 5.462 | 0.021* |
| Temperature | 0.613 | 0.435 | 0.991 | 0.321 | 1.482 | 0.226 |
| Light Intensity | 1.123 | 0.291 | 3.503 | 0.063 | 1.992 | 0.161 |
| Infection-by-Sex | 0.364 | 0.547 | 7.076 | 0.008* | 0.068 | 0.794 |
| Sample Size | 204 | | 240 | | 147 | |

**Supplemental Table S3.** Results from linear models of square root-transformed mean absolute distance sums (ADS). Leverage and distribution analyses revealed that the mean ADS data for *D. melanogaster*, *D. yakuba*, *D. santomea*, and *D. teissieri* required a square root transformation. Significant effects identified by *F* tests at the *P* = 0.05 threshold are marked with an asterisk.

| | ***w*MelCS** | | ***w*Mel (Maine)** | | ***w*Mel (Panama)** | | ***w*Yak** | | ***w*San** | |
|---|---|---|---|---|---|---|---|---|---|---|
| **Explanatory variable** | ***F*** | ***P* value** | ***F*** | ***P* value** | ***F*** | ***P* value** | ***F*** | ***P* value** | ***F*** | ***P* value** |
| Infection Status | 15.857 | <0.001* | 0.249 | 0.618 | 0.261 | 0.610 | 0.170 | 0.681 | 1.305 | 0.255 |
| Sex | 5.572 | 0.019* | 153.961 | <0.001* | 1.034 | 0.311 | 67.904 | <0.001* | 114.888 | <0.001* |
| Chamber | 3.914 | <0.001* | 5.223 | <0.001* | 1.505 | 0.117 | 3.719 | <0.001* | 1.682 | 0.062 |
| Start Time | 6.642 | 0.010* | 18.839 | <0.001* | 13.984 | <0.001* | 0.393 | 0.532 | 0.178 | 0.674 |
| Water Vapor Pressure | 12.060 | <0.001* | 0.323 | 0.570 | 0.026 | 0.871 | 0.112 | 0.739 | 11.196 | <0.001* |
| Relative Humidity | 0.858 | 0.355 | 4.462 | 0.035* | 1.257 | 0.264 | 0.125 | 0.724 | 1.132 | 0.289 |
| Temperature | 0.252 | 0.616 | 1.326 | 0.250 | 3.959 | 0.049* | 5.032 | 0.026* | 24.250 | <0.001* |
| Light Intensity | 0.654 | 0.419 | 0.508 | 0.476 | 0.448 | 0.505 | 0.981 | 0.324 | 0.262 | 0.610 |
| Infection-by-Sex | 4.566 | 0.033* | 1.247 | 0.265 | 0.113 | 0.737 | 0.066 | 0.797 | 1.051 | 0.307 |
| Sample Size | 464 | | 341 | | 160 | | 164 | | 209 | |

| | ***w*Tei** | | ***w*Ri** | | ***w*Suz** | | ***w*Aur** | | ***w*Ha** | |
|---|---|---|---|---|---|---|---|---|---|---|
| **Explanatory variable** | ***F*** | ***P* value** | ***F*** | ***P* value** | ***F*** | ***P* value** | ***F*** | ***P* value** | ***F*** | ***P* value** |
| Infection Status | 0.661 | 0.417 | 5.671 | 0.019* | 7.155 | 0.008* | 0.690 | 0.408 | 5.112 | 0.024* |
| Sex | 93.461 | <0.001* | 0.091 | 0.763 | 7.676 | 0.006* | 55.467 | <0.001* | 2.377 | 0.124 |
| Chamber | 5.125 | <0.001* | 0.847 | 0.624 | 1.238 | 0.254 | 3.420 | <0.001* | 1.793 | 0.038* |
| Start Time | 0.715 | 0.398 | 2.835 | 0.095 | 3.941 | 0.049* | 0.516 | 0.473 | 0.491 | 0.484 |
| Water Vapor Pressure | 7.491 | 0.007* | 0.822 | 0.367 | 0.775 | 0.380 | 6.383 | 0.013* | 1.005 | 0.317 |
| Relative Humidity | 0.119 | 0.731 | 0.007 | 0.935 | 4.985 | 0.027* | 9.326 | 0.003* | 0.270 | 0.604 |
| Temperature | 3.864 | 0.050* | 0.139 | 0.710 | 8.652 | 0.004* | 0.710 | 0.401 | 0.749 | 0.388 |
| Light Intensity | 6.912 | 0.009* | 3.633 | 0.059 | 7.086 | 0.009* | 11.807 | <0.001* | 0.503 | 0.479 |
| Infection-by-Sex | 1.516 | 0.219 | 7.167 | 0.009* | 0.035 | 0.851 | 2.305 | 0.131 | 0.104 | 0.747 |
| Sample Size | 347 | | 131 | | 163 | | 181 | | 353 | |

| | ***w*Sh** | | ***w*Ha & *w*No** | | ***w*Mau** | |
|---|---|---|---|---|---|---|
| **Explanatory variable** | ***F*** | ***P* value** | ***F*** | ***P* value** | ***F*** | ***P* value** |
| Infection Status | 0.172 | 0.679 | 10.405 | 0.001* | 0.492 | 0.484 |
| Sex | 9.438 | 0.002* | 43.662 | <0.001* | 0.501 | 0.481 |
| Chamber | 1.733 | 0.053 | 1.462 | 0.127 | 2.775 | 0.001* |
| Start Time | 2.079 | 0.151 | 1.668 | 0.198 | 0.317 | 0.574 |
| Water Vapor Pressure | 2.741 | 0.100 | 1.335 | 0.249 | 0.573 | 0.451 |
| Relative Humidity | 0.282 | 0.596 | 10.193 | 0.002* | 4.102 | 0.045* |
| Temperature | 0.226 | 0.635 | 3.190 | 0.075 | 2.573 | 0.111 |
| Light Intensity | 0.395 | 0.530 | 2.633 | 0.106 | 0.335 | 0.564 |
| Infection-by-Sex | 1.188 | 0.277 | 9.136 | 0.003* | 0.038 | 0.845 |
| Sample Size | 204 | | 240 | | 147 | |

**Supplemental Table S4.** Results from linear models of log-transformed "activity events" count data, the number of seconds that each fly triggered the infrared activity monitor over the 3-hour period. Leverage and distribution analyses revealed that the count data for *D. simulans*, *D. suzukii, D. auraria*, *D. mauritiana*, and *D. sechellia* required a log transformation. Significant effects identified by *F* tests at the *P* = 0.05 threshold are marked with an asterisk.

| | ***w*MelCS** | | ***w*Mel (Maine)** | | ***w*Mel (Panama)** | | ***w*Yak** | | ***w*San** | |
|---|---|---|---|---|---|---|---|---|---|---|
| **Explanatory variable** | ***F*** | ***P* value** | ***F*** | ***P* value** | ***F*** | ***P* value** | ***F*** | ***P* value** | ***F*** | ***P* value** |
| Infection Status | 11.872 | <0.001* | 1.870 | 0.172 | 0.997 | 0.320 | 0.009 | 0.924 | 0.800 | 0.372 |
| Sex | 16.614 | <0.001* | 131.112 | <0.001* | 2.336 | 0.129 | 69.517 | <0.001* | 113.588 | <0.001* |
| Chamber | 0.970 | 0.483 | 1.931 | 0.023* | 1.263 | 0.239 | 1.189 | 0.290 | 1.203 | 0.276 |
| Start Time | 5.371 | 0.021* | 20.839 | <0.001* | 11.311 | <0.001* | 0.218 | 0.641 | 0.098 | 0.755 |
| Water Vapor Pressure | 6.001 | 0.015* | 0.256 | 0.613 | 1.043 | 0.309 | 0.376 | 0.541 | 13.753 | <0.001* |
| Relative Humidity | 1.867 | 0.172 | 3.642 | 0.057 | 0.676 | 0.412 | 1.239 | 0.268 | 1.800 | 0.181 |
| Temperature | 1.581 | 0.209 | 1.180 | 0.278 | 5.248 | 0.024* | 2.727 | 0.101 | 26.949 | <0.001* |
| Light Intensity | 0.493 | 0.483 | 0.229 | 0.632 | 0.001 | 0.980 | 0.057 | 0.813 | 0.342 | 0.559 |
| Infection-by-Sex | 3.440 | 0.064 | 2.552 | 0.111 | 0.231 | 0.632 | 0.046 | 0.831 | 0.405 | 0.525 |
| Sample Size | 464 | | 341 | | 160 | | 164 | | 209 | |

| | ***w*Tei** | | ***w*Ri** | | ***w*Suz** | | ***w*Aur** | | ***w*Ha** | |
|---|---|---|---|---|---|---|---|---|---|---|
| **Explanatory variable** | ***F*** | ***P* value** | ***F*** | ***P* value** | ***F*** | ***P* value** | ***F*** | ***P* value** | ***F*** | ***P* value** |
| Infection Status | 0.410 | 0.522 | 4.197 | 0.043* | 8.307 | 0.005* | 1.389 | 0.240 | 6.952 | 0.009* |
| Sex | 85.186 | <0.001* | 0.411 | 0.523 | 3.030 | 0.084 | 78.024 | <0.001* | 4.822 | 0.029* |
| Chamber | 2.570 | 0.002* | 0.959 | 0.504 | 1.933 | 0.028* | 2.087 | 0.015* | 4.460 | <0.001* |
| Start Time | 4.690 | 0.031* | 4.431 | 0.038 | 6.653 | 0.011* | 1.292 | 0.257 | 2.859 | 0.092 |
| Water Vapor Pressure | 15.686 | <0.001* | 0.001 | 0.982 | 1.299 | 0.256 | 23.205 | <0.001* | 1.646 | 0.200 |
| Relative Humidity | 0.013 | 0.909 | 0.420 | 0.518 | 3.970 | 0.048* | 7.553 | 0.007* | 0.404 | 0.525 |
| Temperature | 1.222 | 0.270 | 0.705 | 0.403 | 8.975 | 0.003* | 0.249 | 0.619 | 1.344 | 0.247 |
| Light Intensity | 3.617 | 0.058 | 2.836 | 0.095 | 11.039 | 0.001* | 21.524 | <0.001* | 0.821 | 0.366 |
| Infection-by-Sex | 0.598 | 0.440 | 8.437 | 0.004* | 0.019 | 0.891 | 7.621 | 0.006* | 0.421 | 0.517 |
| Sample Size | 347 | | 131 | | 163 | | 181 | | 353 | |

| | ***w*Sh** | | ***w*Ha & *w*No** | | ***w*Mau** | |
|---|---|---|---|---|---|---|
| **Explanatory variable** | ***F*** | ***P* value** | ***F*** | ***P* value** | ***F*** | ***P* value** |
| Infection Status | 0.924 | 0.338 | 5.528 | 0.020* | 0.010 | 0.919 |
| Sex | 23.014 | <0.001* | 33.568 | <0.001* | 0.038 | 0.847 |
| Chamber | 3.127 | <0.001* | 1.099 | 0.360 | 4.670 | <0.001* |
| Start Time | 1.088 | 0.298 | 4.444 | 0.036* | 1.416 | 0.236 |
| Water Vapor Pressure | 8.133 | 0.005* | 2.005 | 0.158 | 0.422 | 0.517 |
| Relative Humidity | 0.158 | 0.691 | 11.318 | <0.001* | 6.078 | 0.015* |
| Temperature | 0.258 | 0.612 | 1.475 | 0.226 | 2.366 | 0.127 |
| Light Intensity | 0.418 | 0.519 | 2.150 | 0.144 | 1.334 | 0.250 |
| Infection-by-Sex | 0.012 | 0.914 | 6.336 | 0.013* | 0.211 | 0.647 |
| Sample Size | 204 | | 240 | | 147 | |

**Supplemental Table S5.** Results from linear models of square root-transformed "activity events" count data, the number of seconds that each fly triggered the infrared activity monitor over the 3-hour period. Leverage and distribution analyses revealed that the count data for *D. melanogaster*, *D. yakuba*, *D. santomea*, and *D. teissieri* required a square root transformation. Significant effects identified by *F* tests at the *P* = 0.05 threshold are marked with an asterisk.

| | ***w*MelCS** | | ***w*Mel (Maine)** | | ***w*Mel (Panama)** | | ***w*Yak** | | ***w*San** | |
|---|---|---|---|---|---|---|---|---|---|---|
| **Explanatory variable** | ***F*** | ***P* value** | ***F*** | ***P* value** | ***F*** | ***P* value** | ***F*** | ***P* value** | ***F*** | ***P* value** |
| Infection Status | 14.735 | <0.001* | 0.449 | 0.503 | 0.425 | 0.516 | 0.038 | 0.846 | 1.997 | 0.159 |
| Sex | 12.062 | <0.001* | 154.420 | <0.001* | 0.879 | 0.350 | 76.558 | <0.001* | 124.819 | <0.001* |
| Chamber | 1.707 | 0.051 | 2.784 | <0.001* | 1.193 | 0.287 | 1.975 | 0.024* | 1.039 | 0.417 |
| Start Time | 7.068 | 0.008* | 19.541 | <0.001* | 14.292 | <0.001* | 0.008 | 0.928 | 0.034 | 0.853 |
| Water Vapor Pressure | 7.965 | 0.005* | 0.597 | 0.440 | 0.111 | 0.739 | 0.054 | 0.817 | 10.585 | 0.001* |
| Relative Humidity | 1.017 | 0.314 | 4.433 | 0.036* | 2.046 | 0.155 | 0.550 | 0.459 | 1.176 | 0.280 |
| Temperature | 0.721 | 0.396 | 1.183 | 0.278 | 3.934 | 0.049* | 3.872 | 0.051 | 24.471 | <0.001* |
| Light Intensity | 0.271 | 0.603 | 0.033 | 0.856 | 0.164 | 0.686 | 0.213 | 0.646 | 0.608 | 0.436 |
| Infection-by-Sex | 4.637 | 0.032* | 2.211 | 0.138 | 0.164 | 0.686 | 0.000 | 0.984 | 1.440 | 0.232 |
| Sample Size | 464 | | 341 | | 160 | | 164 | | 209 | |

| | ***w*Tei** | | ***w*Ri** | | ***w*Suz** | | ***w*Aur** | | ***w*Ha** | |
|---|---|---|---|---|---|---|---|---|---|---|
| **Explanatory variable** | ***F*** | ***P* value** | ***F*** | ***P* value** | ***F*** | ***P* value** | ***F*** | ***P* value** | ***F*** | ***P* value** |
| Infection Status | 0.603 | 0.438 | 5.764 | 0.018* | 7.114 | 0.009* | 0.522 | 0.471 | 4.757 | 0.030* |
| Sex | 106.138 | <0.001* | 0.195 | 0.659 | 7.582 | 0.007* | 78.838 | <0.001* | 2.909 | 0.089 |
| Chamber | 3.857 | <0.001* | 0.990 | 0.471 | 1.426 | 0.148 | 2.723 | 0.001* | 2.238 | 0.007* |
| Start Time | 1.424 | 0.234 | 3.387 | 0.068 | 3.831 | 0.052 | 1.402 | 0.238 | 0.735 | 0.392 |
| Water Vapor Pressure | 15.920 | <0.001* | 0.796 | 0.374 | 0.473 | 0.493 | 16.036 | <0.001* | 1.870 | 0.172 |
| Relative Humidity | 0.042 | 0.837 | 0.000 | 0.999 | 5.771 | 0.018* | 7.937 | 0.005* | 0.370 | 0.543 |
| Temperature | 3.414 | 0.066 | 0.061 | 0.805 | 9.504 | 0.002* | 0.003 | 0.959 | 0.775 | 0.379 |
| Light Intensity | 5.889 | 0.016* | 4.007 | 0.048* | 7.563 | 0.007* | 18.003 | <0.001* | 0.568 | 0.452 |
| Infection-by-Sex | 1.106 | 0.294 | 7.036 | 0.009* | 0.123 | 0.726 | 4.332 | 0.039* | 0.205 | 0.651 |
| Sample Size | 347 | | 131 | | 163 | | 181 | | 353 | |

| | ***w*Sh** | | ***w*Ha & *w*No** | | ***w*Mau** | |
|---|---|---|---|---|---|---|
| **Explanatory variable** | ***F*** | ***P* value** | ***F*** | ***P* value** | ***F*** | ***P* value** |
| Infection Status | 0.762 | 0.384 | 10.326 | 0.002* | 0.360 | 0.550 |
| Sex | 12.470 | <0.001* | 43.969 | <0.001* | 0.092 | 0.763 |
| Chamber | 2.190 | 0.010* | 1.171 | 0.299 | 4.530 | <0.001* |
| Start Time | 1.989 | 0.160 | 2.328 | 0.129 | 0.613 | 0.435 |
| Water Vapor Pressure | 5.668 | 0.018* | 1.105 | 0.294 | 1.922 | 0.168 |
| Relative Humidity | 0.214 | 0.644 | 10.004 | 0.002* | 6.071 | 0.015* |
| Temperature | 0.118 | 0.732 | 3.288 | 0.071 | 4.525 | 0.035* |
| Light Intensity | 0.335 | 0.563 | 1.803 | 0.181 | 0.000 | 0.989 |
| Infection-by-Sex | 0.320 | 0.572 | 7.852 | 0.006* | 0.025 | 0.875 |
| Sample Size | 204 | | 240 | | 147 | |

**Supplemental Table S6.** The scaffold count, N50, and total assembly size of each *Wolbachia* assembly.

| Genome | Host Genotype ID | Scaffold Count | N50 | Total Assembly Size |
|---|---|---|---|---|
| *w*Aur | *SP11-11* (Accession PRJNA432099) | 116 | 20,018 | 1,319,217 |
| *w*Mel (Maine) | *FFD25* | 89 | 25,061 | 1,241,721 |

### SUPPLEMENTAL FIGURES

**Supplemental Figure S1.** Locomotor activity was measured in a 16-chamber flow-through respirometry and data acquisition system (MAVEn, Sable Systems International). The MAVEn includes 16 2.4 ml volume polycarbonate animal chambers and an activity board that uses infrared light (invisible to flies) to monitor animal activity in each chamber, sampled at 1 Hz (shown on the left). The activity board also includes sensors that record ambient temperature, humidity, and light intensity within the measurement area. For activity assays, a constant stream of dry, $CO_2$-free air from a compressed air cylinder was humidified by flowing through Nafion tubing submerged in distilled water and then directed into each chamber at a flow rate of 35 ml/min. All assays were conducted at room temperature (22.9°C ± 0.01 s.e.). Chambers were washed with warm soapy water between runs. Prior to each activity assay, a white acrylic reflecting cover was positioned on top of the chambers (shown on the right) to secure the chambers above the activity monitors, block ambient lab light, and optimize infrared reflection off the flies.

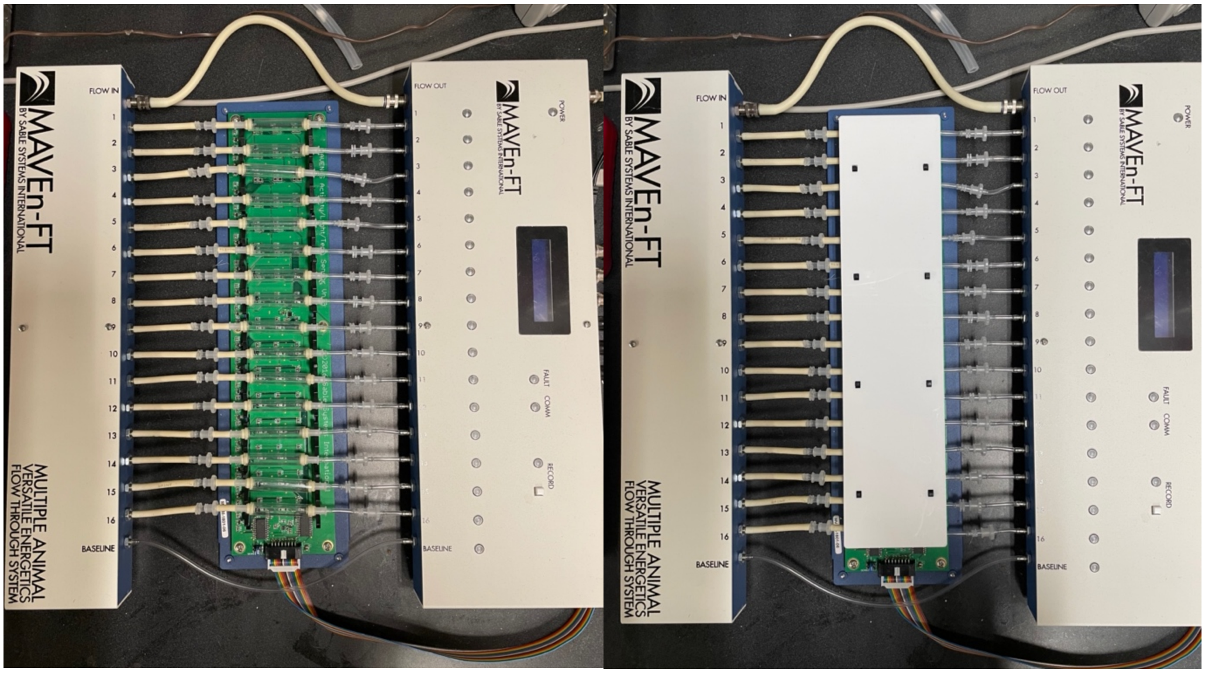

**Supplemental Figure S2.** Results from activity assays when activity is measured as the number of "activity events" or the number of seconds that each fly triggered the infrared activity monitor over the 3-hour period. Significance was evaluated using linear models (Tables S4 and S5).

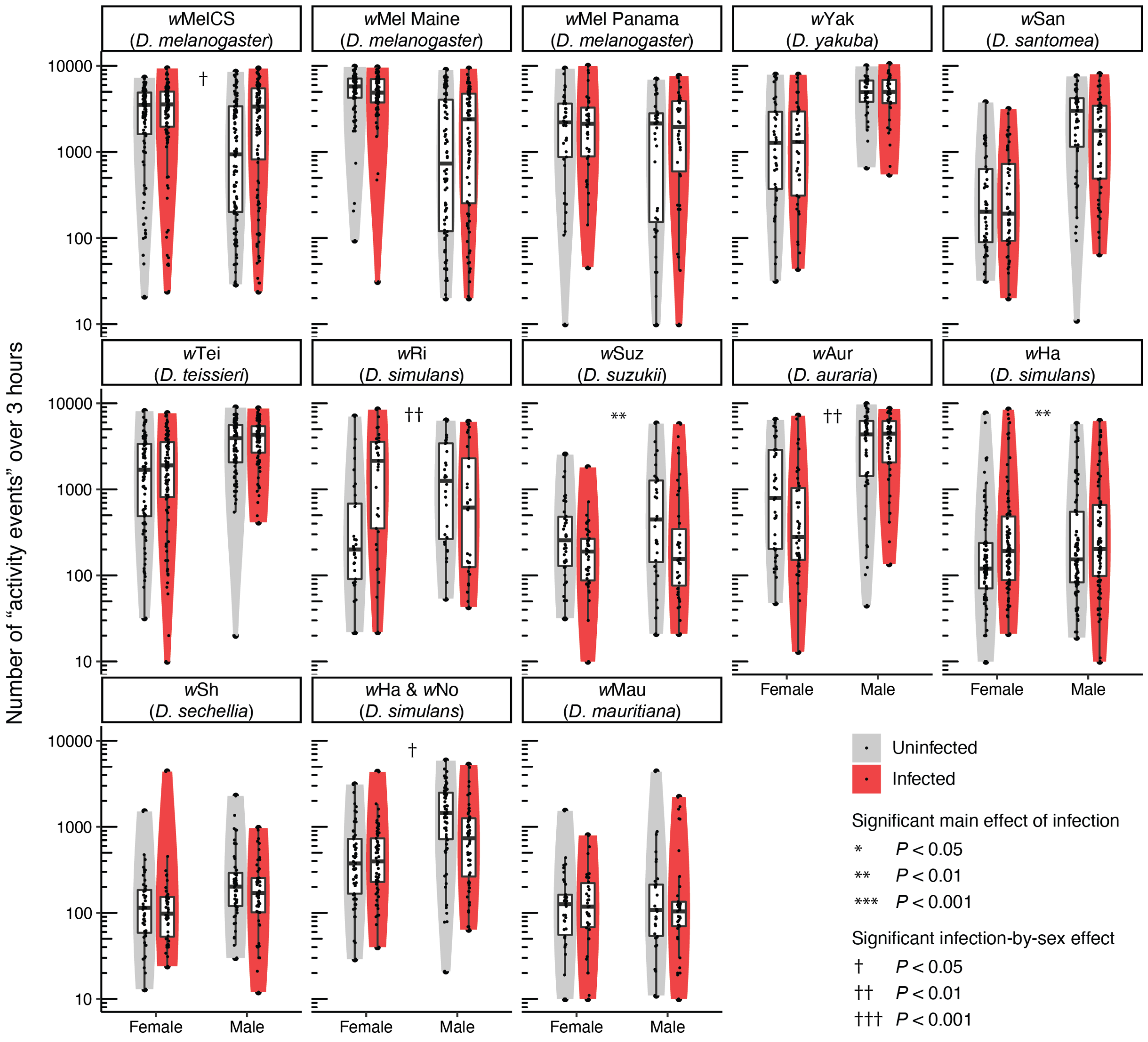

**Supplemental Figure S3.** Estimated Bayesian phylogram for A- and B-group *Wolbachia* strains examined in this study. The phylogram was estimated with 211 single-copy genes of identical length in all of the genomes, spanning 178,569 bp. All nodes have Bayesian posterior probabilities of 1. The divergence time estimate (million years ago [MYA]) for A- and B-group *Wolbachia* is superimposed from Meany et al. (2019). To the right, the change in least-square (LS) mean square root-transformed activity (ADS) for each sex. LS means were generated from linear models (Table S3).

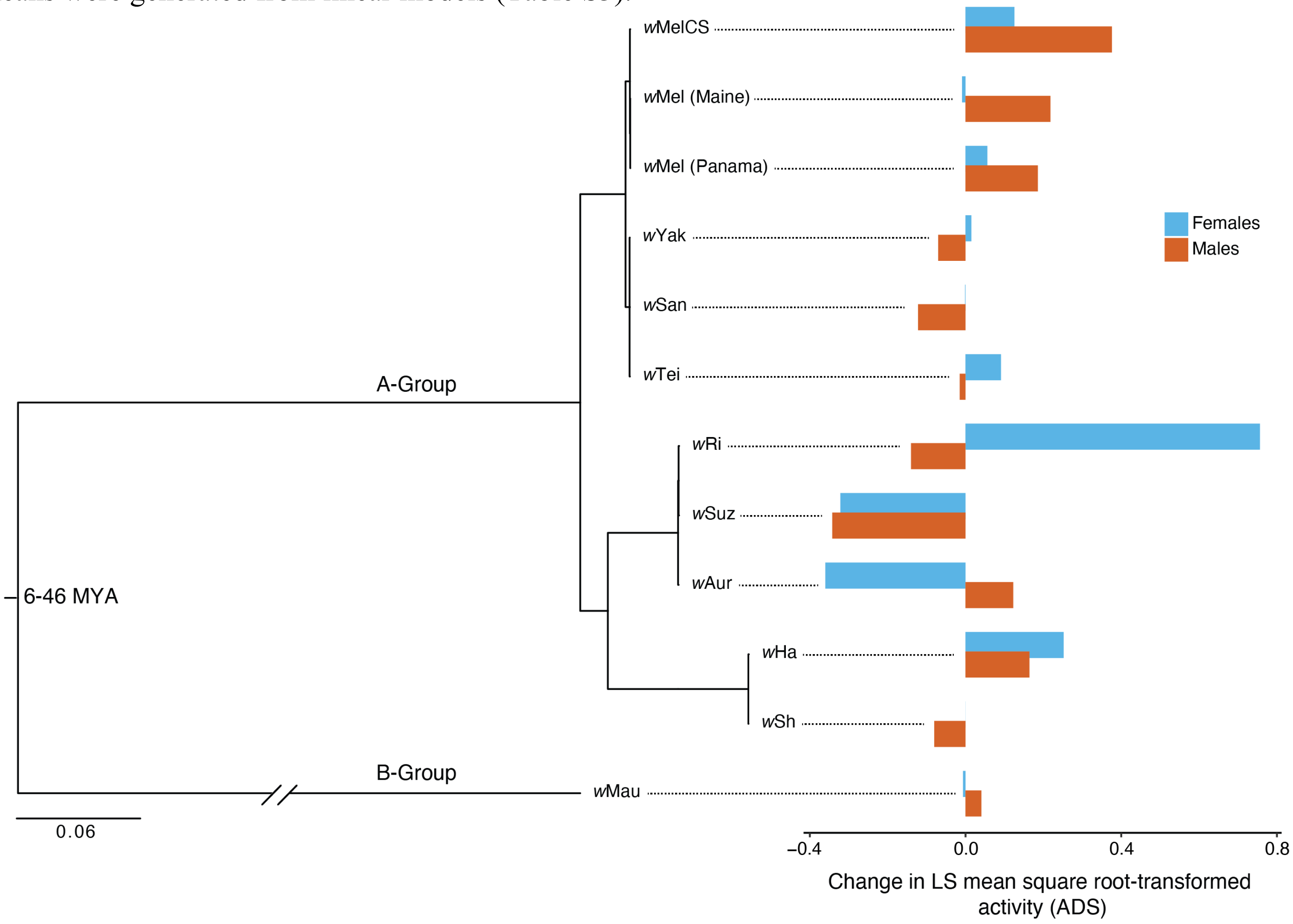

**Supplemental Figure S4.** *D* statistic simulations testing whether *Wolbachia* effects on host activity exhibit phylogenetic signal when treated as a binary trait. The observed *D* statistic is compared to alternative *D* values generated with simulated data based on a random phylogenetic pattern and Brownian motion, using 1,000 permutations each. $D = 1$ is consistent with random trait evolution, whereas $D = 0$ supports a Brownian motion model of character evolution. The *D* statistic is based on the sum of the character changes along the edges of the phylogeny. Above, the scaling of *D* values is shown based on the distributions of simulated sums of character changes under models of random association (blue) and Brownian motion (red). Continuous vertical lines represent the mean of simulated *D* values for phylogenetic randomness (blue) and Brownian motion (red). The dashed line indicates our observed *D* statistic. The large area of overlap between the distributions of simulations representing phylogenetic randomness and Brownian motion suggests it is difficult to distinguish between these two alterative scenarios for a phylogeny of this size ($N = 12$ *Wolbachia* strains).

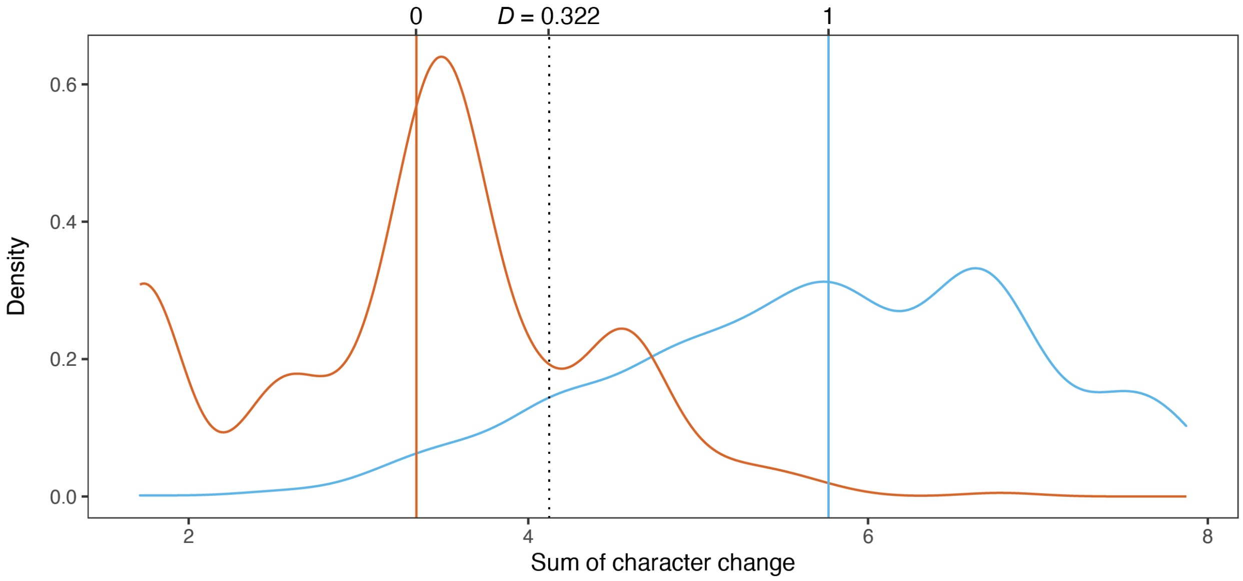

**Supplemental Figure S5.** Mean *D* and 95% confidence intervals generated from 100 replicates of simulated binary trait evolution using an increasing number of *Wolbachia* strains ($N = 12$, 25, 50, 100). We generated simulated trees using the "sim.bdtree" and "sim.char" functions in the *geiger* R package (Harmon et al. 2008). For each tree, we simulated character evolution of a continuous trait with no phylogenetic signal ($\lambda = 0$) and strong signal ($\lambda = 1$). Following Fritz and Purvis (2010), we used a threshold to convert continuous characters to binary traits using a prevalence of 0.417, the proportion of *Wolbachia* strains from our original phylogeny that significantly altered host activity. We then calculated *D* for each tree. The solid horizontal lines show the expectations for *D* using $\lambda = 0$ ($D = 1$, i.e. no phylogenetic signal) and $\lambda = 1$ ($D = 0$; i.e. strong phylogenetic signal). The dotted horizontal line shows our *D* value calculated from the original phylogeny. The simulations show that estimates of *D* are highly variable for small trees, including for trees with the same number of strains as our phylogeny ($N = 12$). As the number of strains increases, estimates of *D* tend to cluster closely around 1 for phylogenetically random traits and around 0 for traits with strong phylogenetic signal. Accordingly, larger trees ($N = 50$, 100) are required to reject a null model of phylogenetic randomness ($D = 1$) at our observed *D* value of 0.322.

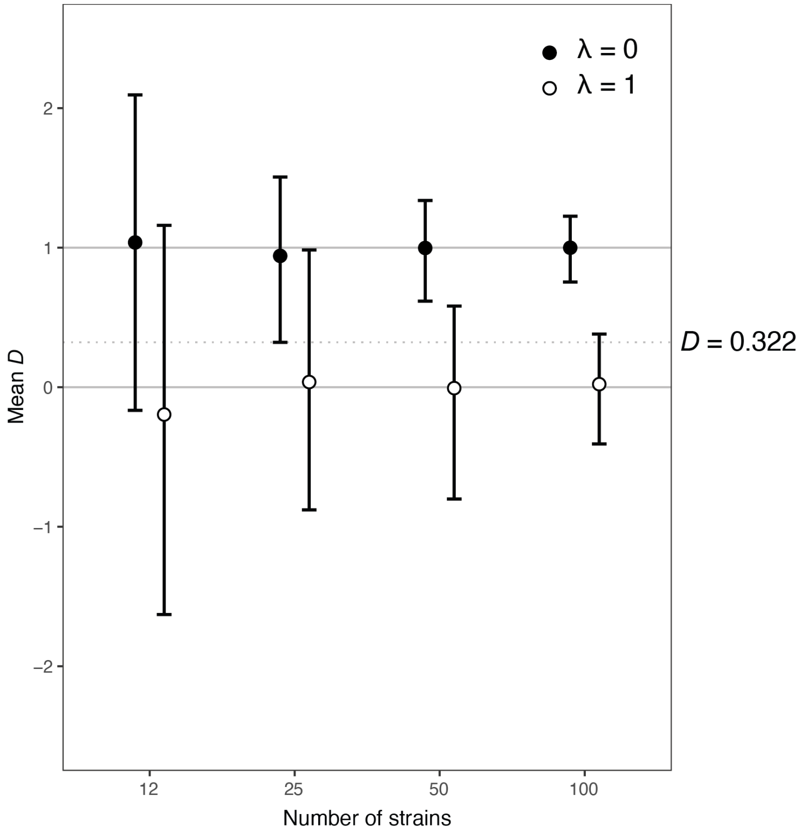

### SUPPLEMETNAL REFERENCES

Ballard, J., and R. Melvin. 2007. Tetracycline treatment influences mitochondrial metabolism and mtDNA density two generations after treatment in *Drosophila*. Insect Molecular Biology 16:799–802.

Chiu, J. C., K. H. Low, D. H. Pike, E. Yildirim, and I. Edery. 2010. Assaying locomotor activity to study circadian rhythms and sleep parameters in *Drosophila*. JoVE (Journal of Visualized Experiments) e2157.

Cooper, B. S., D. Vanderpool, W. R. Conner, D. R. Matute, and M. Turelli. 2019. *Wolbachia* acquisition by *Drosophila yakuba*-clade hosts and transfer of incompatibility loci between distantly related *Wolbachia*. Genetics 212:1399–1419.

Ellegaard, K. M., L. Klasson, K. Näslund, K. Bourtzis, and S. G. Andersson. 2013. Comparative genomics of *Wolbachia* and the bacterial species concept. PLoS Genetics 9.

Fritz, S. A., and A. Purvis. 2010. Selectivity in mammalian extinction risk and threat types: a new measure of phylogenetic signal strength in binary traits. Conservation Biology 24:1042–1051.

Hague, M. T., C. N. Caldwell, and B. S. Cooper. 2020. Pervasive effects of *Wolbachia* on host temperature preference. mBio 11.

Harmon, L. J., J. T. Weir, C. D. Brock, R. E. Glor, and W. Challenger. 2008. GEIGER: investigating evolutionary radiations. Bioinformatics 24:129–131.

Höhna, S., M. J. Landis, T. A. Heath, B. Boussau, N. Lartillot, B. R. Moore, J. P. Huelsenbeck, and F. Ronquist. 2016. RevBayes: Bayesian phylogenetic inference using graphical models and an interactive model-specification language. Systematic Biology 65:726–736.

Jackman, S. D., B. P. Vandervalk, H. Mohamadi, J. Chu, S. Yeo, S. A. Hammond, G. Jahesh, H. Khan, L. Coombe, R. L. Warren, and others. 2017. ABySS 2.0: resource-efficient assembly of large genomes using a Bloom filter. Genome Research 27:768–777.

Joshi, N., J. Fass, and others. 2011. Sickle: A sliding-window, adaptive, quality-based trimming tool for FastQ files.

Klasson, L., J. Westberg, P. Sapountzis, K. Näslund, Y. Lutnaes, A. C. Darby, Z. Veneti, L. Chen, H. R. Braig, R. Garrett, and others. 2009. The mosaic genome structure of the *Wolbachia w*Ri strain infecting *Drosophila simulans*. Proceedings of the National Academy of Sciences 106:5725–5730.

Meany, M. K., W. R. Conner, S. V. Richter, J. A. Bailey, M. Turelli, and B. S. Cooper. 2019. Loss of cytoplasmic incompatibility and minimal fecundity effects explain relatively low *Wolbachia* frequencies in *Drosophila mauritiana*. Evolution 73:1278–1295.

Pfeiffenberger, C., B. C. Lear, K. P. Keegan, and R. Allada. 2010. Locomotor activity level monitoring using the *Drosophila* Activity Monitoring (DAM) System. Cold Spring Harbor Protocols 2010:pdb-prot5518. Cold Spring Harbor Laboratory Press.

Seemann, T. 2014. Prokka: rapid prokaryotic genome annotation. Bioinformatics 30:2068–2069.

Siozios, S., A. Cestaro, R. Kaur, I. Pertot, O. Rota-Stabelli, and G. Anfora. 2013. Draft genome sequence of the *Wolbachia* endosymbiont of *Drosophila suzukii*. Genome announcements 1:e00032-13.

Turelli, M., B. S. Cooper, K. M. Richardson, P. S. Ginsberg, B. Peckenpaugh, C. X. Antelope, K. J. Kim, M. R. May, A. Abrieux, D. A. Wilson, M. J. Bronski, B. R. Moore, J.-J. Gao, M. B. Eisen, J. C. Chiu, W. R. Conner, and A. A. Hoffmann. 2018. Rapid global spread of *w*Ri-like *Wolbachia* across multiple *Drosophila*. Current Biology 28:963-971.e8.
